# Learning to segment self-generated from externally caused optic flow through sensorimotor mismatch circuits

**DOI:** 10.1101/2023.11.15.567170

**Authors:** Matthias Brucklacher, Giovanni Pezzulo, Francesco Mannella, Gaspare Galati, Cyriel M. A. Pennartz

## Abstract

Efficient sensory detection requires the capacity to ignore task-irrelevant information, for example when optic flow patterns created by egomotion need to be disentangled from object perception. To investigate how this is achieved in the visual system, predictive coding with sensorimotor mismatch detection is an attractive starting point. Indeed, experimental evidence for sensorimotor mismatch signals in early visual areas exists, but it is not understood how they are integrated into cortical networks that perform input segmentation and categorization. Our model advances a biologically plausible solution by extending predictive coding models with the ability to distinguish self-generated from externally caused optic flow. We first show that a simple three neuron circuit produces experience-dependent sensorimotor mismatch responses, in agreement with calcium imaging data from mice. This microcircuit is then integrated into a neural network with two generative streams. The motor-to-visual stream consists of parallel microcircuits between motor and visual areas and learns to spatially predict optic flow resulting from self-motion. The second stream bidirectionally connects a motion-selective higher visual area (mHVA) to V1, assigning a crucial role to the abundant feedback connections: the maintenance of a generative model of externally caused optic flow. In the model, area mHVA learns to segment moving objects from the background, and facilitates object categorization. Based on shared neurocomputational principles across species, the model also maps onto primate vision. Our work extends the Hebbian predictive coding to sensorimotor settings, in which the agent actively moves - and learns to predict the consequences of its own movements.

**Significance statement:** This research addresses a fundamental challenge in sensory perception: how the brain distinguishes between self-generated and externally caused visual motion. Using a computational model inspired by predictive coding and sensorimotor mismatch detection, the study proposes a biologically plausible solution. The model incorporates a neural microcircuit that generates sensorimotor mismatch responses, aligning with experimental data from mice. This microcircuit is integrated into a neural network with two streams: one predicting self-motion-induced optic flow and another maintaining a generative model for externally caused optic flow. The research advances our understanding of how the brain segments visual input into object and background, shedding light on the neural mechanisms underlying perception and categorization not only in rodents, but also in primates.

## 1 Introduction

Efficient sensory detection requires the capacity to ignore task-irrelevant information. In visual object perception for example, optic flow patterns generated by external objects are more informative than those resulting from self-motion (Gibson, 1950). Consequently, discerning changes in visual input caused by self-movement versus external factors is a crucial challenge for the visual system (see Fig. 1). In computer vision, optic flow finds frequent application, particularly in figure-ground or object segmentation paradigms (see Anthwal and Ganotra, 2019). Indeed, motion has long been recognized to be a powerful Gestalt cue for distinguishing visual objects from the background (Wertheimer, 1923).

**Figure 1.**
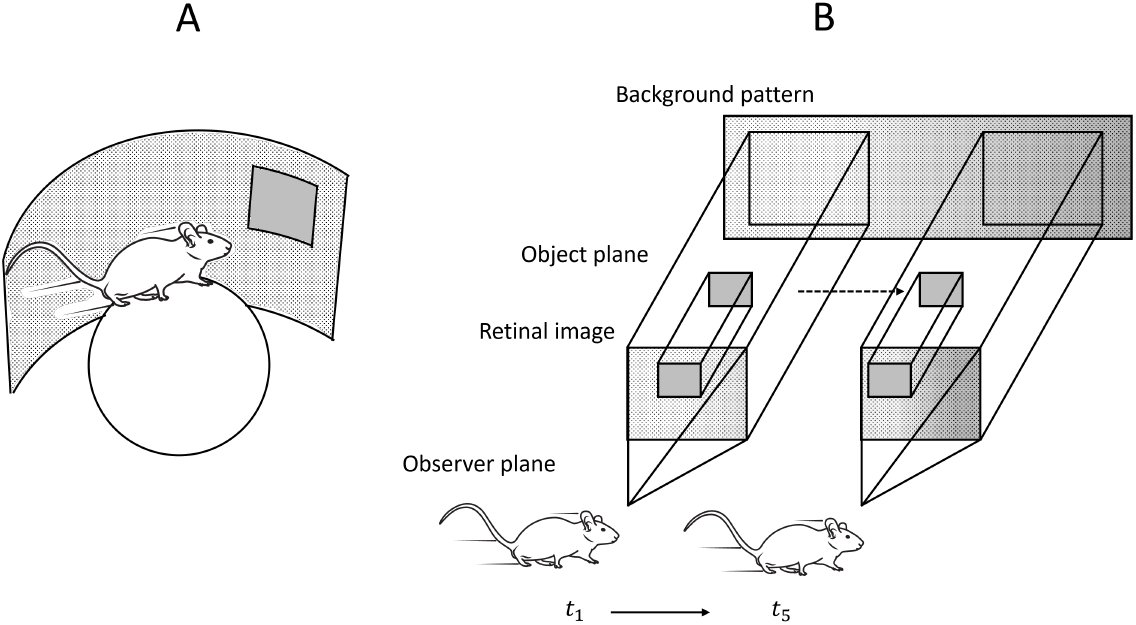
Two causes of sensed optic flow: self-induced and externally generated. (A) Experimental setup for mismatch detection as used in Zmarz and Keller, 2016. The VR environment allows control over the visual input of the mouse as it is moving on a spheric treadmill. (B) Configuration of our simulations, mirroring the image construction process in A. The apparent movement of the background pattern is anticipated based on the motor state of the model. Deviations in optic flow arise when an object moves independently between the background and the observer. Time points *t*_1_ and *t*_5_ depict distinct moments in the simulation.

In the biological context, optic flow has been suggested to be segmented into self- and externally generated components by sensorimotor mismatch - an error signal arising from the disparity between expectation and bottom-up sensory input (Keller & Mrsic-Flogel, 2018). This conjecture gains support from accumulating evidence for the presence of sensorimotor mismatch (or error) signals in early auditory (Audette & Schneider, 2023) and visual areas (Attinger et al., 2017; Zmarz and Keller, 2016, see Fig. 1A). Another recent study on saccadic eye movements by Miura and Scanziani, 2022 demonstrated that direction-selective V1 neurons differentiate between apparent stimulus movement being self- or externally caused.

These findings suggest the presence of a motor-to-sensory forward model conveying corollary discharges from motor areas to early sensory areas and transforming them into sensory coordinates (see Frith et al., 2000 for an overview). Beyond rodents, motor feedback affects sensory activity in primates, too (Eliades & Wang, 2008; Pelah & Barlow, 1996; Zhou et al., 2016). This endows the brain with neural circuits for mismatch (or prediction error)-based segmentation.

While models demonstrating sensorimotor mismatch responses exist (Hertäg & Sprekeler, 2020; Mikulasch et al., 2022), it remains unclear if and how these can functionally contribute to segmentation operations. Moreover, given that a significant portion of visual input is generated by external motion, inferring the causes of optic flow becomes a two-fold problem. This raises the question how predictions from the motor-to-sensory forward model and predictions about external objects integrate mechanistically to distinguish between self-caused and externally generated optic flow and interpret them.

Here, we show that the framework of predictive coding elegantly and parsimoniously allows integration of these two generative processes. Predictive coding describes perception as a hierarchical generative process in which higher cortical areas improve their top-down prediction of activity in lower areas (Friston, 2005; Lee & Mumford, 2003; Rao & Ballard, 1999). It forms an important computational building block for understanding how perception – and in particular a motion-corrected, stable world representation - is constructed through learning and inference (Andersen et al., 1985; Crapse & Sommer, 2008; Creutzig & Sprekeler, 2008; Pennartz, 2015; Whishaw & Brooks, 1999). In pure form, both inference (updating neural activities) and learning of synaptic weights are driven by prediction errors. In contrast to most predictive coding models derived from Rao and Ballard, 1999, however, our model does not predict luminosity, but optic flow. First, a core microcircuit is constructed, with its biological plausibility strengthened by replication of experimentally observed sensorimotor mismatch (Attinger et al., 2017). Second, the model is scaled up and extended by circuits for the representation of externally generated optic flow, leading to segmentation of self-vs. externally generated inputs (Fig. 1B). Lastly, we demonstrate that the motion-selective higher visual areas of the model, become tuned to optic flow patterns caused by external objects, facilitating classification of the perceived object. The model proposed here thus extends biologically plausible predictive coding, by functionally integrating motor-to-visual feedback signals in multi-area generative inference of optic flow.

## 2 Methods

### 2.1 A microcircuit for sensorimotor mismatch detection

We developed a microcircuit for sensorimotor mismatch calculation that is shown in Fig. 2A. In inference, optic flow elicits neural activity *y*_*vis*_ ∈ [0, 1] of a direction selective cell in V1 (as originally observed in cats by Hubel and Wiesel, 1959). Simultaneously, the motor area (roughly corresponding to motor cortex or closely connected areas (Guitchounts et al., 2020; Leinweber et al., 2017), here represented by only one unit) codes for the current motor state via neural activity *y*_*mot*_ ∈ [0, 1] of a subpopulation. This motor state is transformed to sensory coordinates via a forward model (Frith et al., 2000).

**Figure 2.**
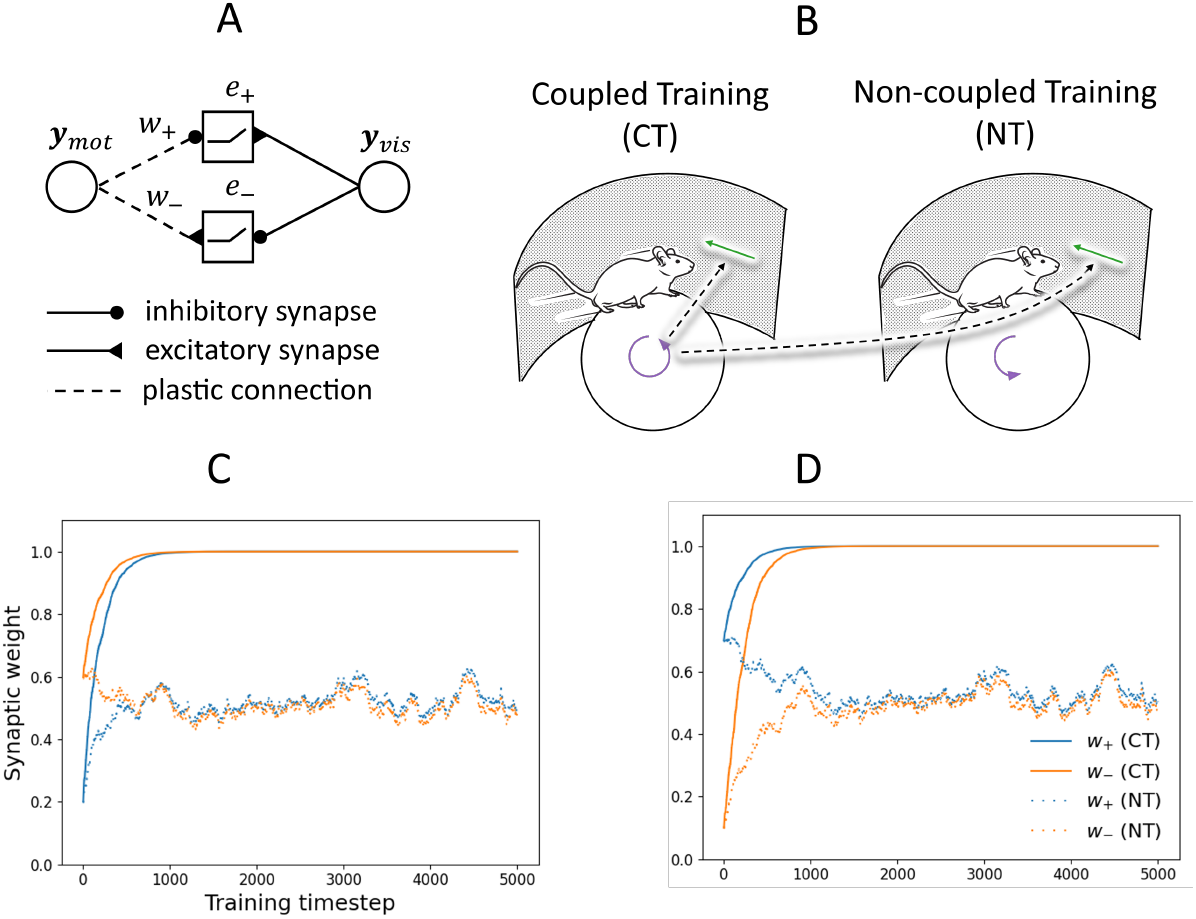
Proposed microcircuit for mismatch calculation and weight evolution during training. (A) Proposed microcircuit with error neurons depicted as squares and representation neurons as circles. *w*_−_ is the weight of the forward model’s excitatory connection to the negative prediction error neuron, *w*_+_ the weight of the inhibitory connection to the positive prediction error neuron (see Equation 1. The predicted optic flow is given as the outcome of the forward model, i.e. the synaptic currents arriving at the error neurons from the left. (B) Illustration of coupled and non-coupled training: in both cases, visual inputs are the same, given by the motor movements in the CT condition. In NT, however, motor states and visual inputs are completely uncorrelated. (C) Illustration of robustness to initial conditions proven in section 7.1: Shown is the synaptic weight in coupled (CT) and non-coupled (NT) training (see main text) over training time. (D) Same as (C) for a different weight initialization scheme.

Inversely connected via excitatory and inhibitory synapses, positive and negative error neurons then compare the sensory input to the predictions from the forward model. We chose to model positive and negative error neurons separately, as they are considered to offer a better fit to experimental data (Keller & Mrsic-Flogel, 2018). Their firing rate is given as

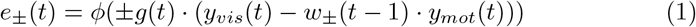

(note the inverse wiring of positive and negative error neurons illustrated in Figure 2A), with synaptic weights *w*_*±*_ of the forward model (determined by learning at the previous time step described below or in the initial condition), gating function *g*(*t*) that keeps the firing rate at baseline if the model is not sending a movement signal (*y*_*mot*_ = 0). This gating is putatively mediated by signalling from higher-order thalamus or neuromodulation (Keller & Mrsic-Flogel, 2018):

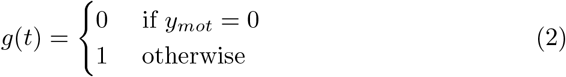

and activation function

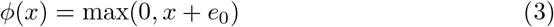

with baseline firing rate *e*_0_ = 0.5 and the total synaptic input *x*. Supportive evidence for bipartite error computation comes from biologically detailed models that show how these circuits can develop in cortical tissue (Hertäg & Sprekeler, 2020; Mirasso et al., 2023). Learning is then mediated by a Hebbian rule with a switch between long-term potentiation and long-term depression mediated by NMDA receptor activation of the postsynapse (Lüscher & Malenka, 2012; Malenka, 1994):

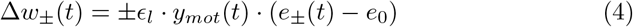

with learning rate *ϵ*_*l*_ = 0.01. In supplementary section 7.1, a proof of convergence of this learning rule is provided.

For comparison with Attinger et al., 2017, where two cohorts of mice were raised differently, under controlled regimes of sensorimotor experience, we trained two copies of the microcircuit with input contingencies based on their paradigm. Importantly, both visual input and motor state are one-dimensional at this point. In coupled training (CT), the motor state of the animal, sampled from a Bernoulli distribution, is reflected in the optic flow: *y*_*vis*_ = *y*_*mot*_ ∼*ℬ* (1, 0.5). In the non-coupled training paradigm (NT), motor states are the same as in the CT condition, but optic flow is independently sampled from an identical distribution. In both conditions, convergence of the forward model weights was robust to different weight initialization and depended strongly on the training paradigm (CT or NT; Fig. 2B-C).

### 2.2 Integration with hierarchical predictive coding

#### 2.2.1 Inference in the two-stream architecture

The presence of mismatch computation in primary visual cortex fits well with theories of generative perception in which the brain interprets visual inputs by identifying the causes that are most likely to have produced them (Dayan et al., 1995; Friston, 2005; Gregory, 1980; Lee & Mumford, 2003; Pennartz, 2015; Rao & Ballard, 1999; von Helmholtz, 1867). We thus integrated the microcircuit developed above into a well-known framework of modeling generative perception, hierarchical predictive coding, which can in principle be applied to rodent and primate visual cortex. While functional and hierarchical modularization is well known in primate visual cortex, also the mouse homolog is hierarchically structured and functionally modularized (Andermann et al., 2011; Marshel et al., 2011; Siegle et al., 2021; Wang et al., 2011).

Visual inputs to the model are multidimensional (we used a retinotopic field of 40 x 40 or 80 x 80 units depending on the dataset used), covering the visual field. As in most predictive coding models, updating of neural activities (inference) and updating of synaptic strengths (learning) are split into separate phases. During inference, the model receives grayscale video as input. From this, optic flow is extracted through the Farnebäck algorithm (Farnebäck, 2003), mimicking the functioning of direction selective cells in V1. As a result, four populations of cells encode orthogonal stimulus movement directions (left/right/up/down) leading to a V1 population of 40 x 40 x 4 = 6,400 neurons (for the modified FashionMNIST dataset) and 80 x 80 x 4 = 25,600 neurons for the Animal dataset respectively). An equal number of positive and negative error neurons then compares velocity of sensory, bottom-up optic flow in the respective direction to top-down predictions, fitting well with reports of retinotopic mismatch calculation (Zmarz & Keller, 2016). These are mediated via two pathways shown in Fig. 3 that jointly attempt to explain the sensed optic flow:

**Figure 3.**
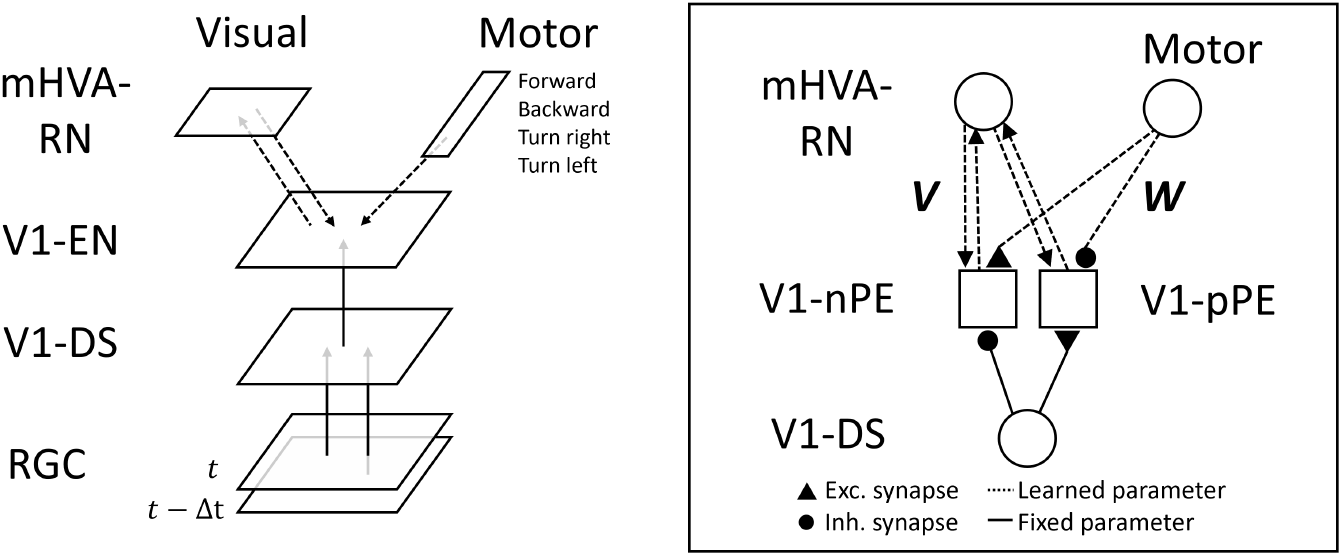
Integration of motor-to-sensory predictions with predictions from higher visual areas. Left: Error neurons in visual area V1 receive predictions about optic flow from two streams. After training, the motor-to-visual stream sends predictions about expected optic flow as a result of egomotion. The remaining prediction errors in V1 are fed forward to the motion-selective higher visual area (mHVA). Right: Synaptic wiring of subpopulations with error neurons depicted as squares and representation neurons as circles. Synaptic connections subject to the Hebbian learning rule of Equation 4 are shown as dashed lines: ***V*** for the visual-to-visual stream and ***W*** for the motor-to-visual stream. Connections between mHVA and V1 are not *a priori* constrained to be excitatory or inhibitory and are thus denoted as arrows.

1. The motor-to-sensory forward model consisting of many microcircuits, as developed in section 2.1. The parallel microcircuits form fully connected layers of synaptic connections from a small number *d*_*mov*_ (’dimensionality of the movement space’) of neurons or subpopulations in the motor area to the nPEs and pPEs in V1. Each of the motor neurons codes for a motor-program (moving forward, turning sideways etc.). We begin with a single motor-program (moving forward) and then later extend to *d*_*mov*_ *>* 1 (see Fig. 10). In this pathway, initial weights are drawn from a Gaussian distribution 𝒩 (0.6, 0.6).
2. The visual-to-visual stream originating from model area mHVA and projecting to primary visual area V1. Based on the high fraction of motion-selective neurons, retinotopic organization and bidirectional connection to V1, model area mHVA is taken to correspond to anterolateral (AL), rostrolateral (RL), anteromedial (AM) and posteromedial (PM) cortex, or a subset of these areas, in mice (Andermann et al., 2011; Marshel et al., 2011).

The updating of neural activity in model area mHVA is based on the inference mechanism of Rao and Ballard, 1999: mHVA representation neurons receive inputs from V1 error neurons through convolution kernels of size 4 x 4 x 4 (stride 1), mimicking receptive fields, and inhibit the error neurons in return (Fişek et al., 2023). We used separate sets of weights connecting to pPEs and nPEs, as well as for the four directions each. Depending on the resolution of the dataset, area mHVA thus contains 77 x 77 x 4 = 23,716 or 37 x 37 x 4x4,356 neurons. Initial weights of the visual-to-visual were sampled from 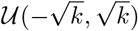where

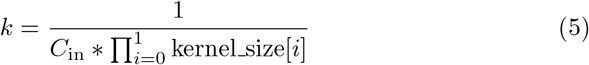

with *C*_*in*_ being the number of feature maps in the higher area and the index *i* enumerating the two spatial dimensions of the quadratic kernel. The equation governing updating of a mHVA neuron’s state variable (akin to the membrane potential) is:

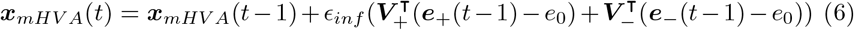

with inference rate *ϵ*_*inf*_ = 0.05, visual weights ***V***_***±***_ from mHVA to V1-EN (cf. Figure 3), and firing rate ***e*** of positive and negative error neurons in V1. ***e***_**0**_ = 0.5 again denotes the baseline firing rate of V1-PE neurons. A ReLU activation function transfers ***x***_*mHV A*_ into the output firing rate

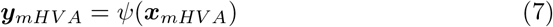

Error neurons in turn have output firing rates determined by (cf. Eq. 1)

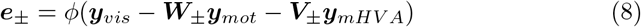

where ***y***_*vis*_ is the magnitude of optic flow in the given direction and ***W***_*±*_ denotes the weights of the motor-to-sensory forward model with motor state ***y***_*mot*_. As previous studies found changes in representational sparsity in deeper layers of unsupervised networks Ororbia and Kifer, 2022, we tested whether another area on top of mHVA (termed mHVA2) showed improved readout accuracy (for details see section 7.6). After inferring neural activity for multiple time steps (15 proved to be sufficient), the Hebbian learning rule from Eq. 4 was used to update synaptic weights.

#### 2.2.2 Training the retinotopic model

Training was conducted in the setting of Fig. 1B, which is comparable to the behavioral paradigm of Attinger et al., 2017; Zmarz and Keller, 2016 (Fig. 1A), but with interpretable objects instead of abstract brick patterns. This paradigm allowed us to study the processing of background and foreground in a controlled manner. However, it also required video inputs with an independently moving object and information about the locomotion state of the observer. As we could not find a dataset of suitable complexity, we constructed three datasets in which movement of the background image is controlled by the model’s motor state:

a. Ten sequences with different animal images moving in front of a background with a grass texture of 80 x 80 pixels (for details see section 7.2). The one-dimensional self-movement signal drives movement of the background patterns across the retina. Sequences consist of blocks with uniform movement speed reflecting locomotion interleaved with blocks of no movement, with a total sequence length of 50 frames. Concerning object motion, several settings are implemented, among others a stable perception paradigm (such as during parallel movement of object and observer, or during gaze following (Zuberbühler, 2008)) and movement of the object image across the retina, independent of self-motion. The dataset also contains a setting with pure background under observer movement and no objects present.
b. Sequences of retinocentric FashionMNIST objects in 40 x 40 resolution with otherwise the same properties as in a), although the FashionMNIST objects are only used in the stable perception paradigm where objects remain retinocentric. Since our modification of this datasets contains a significantly larger number of samples (we use 5000) and is organized in ten object classes, it is suitable to test how useful the neural representations are in a given model area for downstream classification. Across the ten object classes, we use 400 samples for training and 100 different samples per class for testing. Here, only the stable perception paradigm is implemented (objects appear static on the retina).
c. Six different optic flow patterns including non-homogeneous expanding and contracting flow fields without object present. Each flow pattern is associated with one of six dimensions of the motor state encoded in one-hot manner (turning leftwards, moving forward, etc.).

Unless noted differently, the two streams (mHVA to V1 and motor cortex to V1) were trained in the following manner: During pretraining, only the motor-to-visual stream was activated and trained on visual inputs coupled to the model’s motor state. In this phase, no external objects were presented. Then, in the training phase, objects were introduced (with the observer moving along them unless where noted differently) and the visual-to-visual stream was activated. Training proceeded until prediction errors were sufficiently low. To separately analyze learning in the two streams, the weights in the motor-to-visual stream were frozen during training, but see section 7.4 for experiments with joint training.

### 2.3 Segmentation experiments

#### 2.3.1 Obtaining a segmentation mask from the model

Segmentation masks highlighting external objects were obtained from mHVA →V1-EN feedback, based on reports of figure-ground segmentation in V1 through feedback from higher visual areas (Self et al., 2013). Excitatory top-down signals from area mHVA to V1-EN were obtained by matrix multiplication of the top-down synaptic weights ***V***_*±*_ from Equation 6 and mHVA activity *y*_*mHV A*_). The summed up top-down signals were then thresholded at each point in retinotopic space to yield a segmentation mask: Where signals are larger than

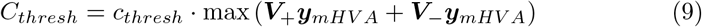

an external object is assumed, with a heuristically chosen scaling constant *c*_*thresh*_. Physiologically, this thresholding may be achieved via a neuron with appropriately set firing threshold via baseline inputs.

#### 2.3.2 Quantifying segmentation accuracy and comparison to baseline

We quantified segmentation accuracy with an Intersection over Union (IoU) measure. IoU is given as the ratio of overlap between predicted and true object area relative to the union and thus ranges between zero and one. Measured at an arbitrarily selected timepoint in the sequence (we chose nine frames into it unless where noted differently), the only important constraint here was object movement relative to the background.

As a strong baseline, we computed a binary segmentation mask ***S***_*baseline*_ by thresholding optic flow signals ***y***_*vis*_:

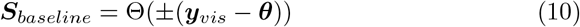

with the Heaviside function Θ: ℝ*→* {0, 1} and threshold ***θ***. The optimal baseline model was then identified by maximizing its IoU score through a grid search of ***θ***.

### 2.4 Linear readout to analyze representational content

Simulating a class-dependent downstream task such as class-specific approach-or-flight behavior, we investigated coding of population activity for object category. Neuronal representations in model areas mHVA, mHVA2, V1 and retina were inferred for training (400 stimuli from each of the ten classes) and test images (100 stimuli per class) from the modified FashionMNIST dataset. Then, a linear classifier (one per each area) was trained to map the activity patterns to class labels using a cross-entropy loss.

## 3 Results

### 3.1 Reproduction of sensorimotor mismatch effects

Before analyzing how well the complete hierarchical network performs computationally, we first examined the biological feasibility of the underlying microcircuit (section 2.1). This involved investigating whether and under what conditions sensorimotor mismatch responses, as discussed in the Introduction, actually appear. Recall that networks were trained in coupled and non-coupled conditions akin to the mice in Attinger et al., 2017. After training, we exposed the networks to the same testing conditions. As in the experimental study, we termed them *mismatch* (continuously moving model, sudden halt of optic flow) and *playback halt* (passive observation of constant optic flow that is suddenly halted). Across both conditions, the error neurons in the model reproduced observed firing rate patterns from mouse V1. As shown in Fig. 4A, a strong increase in neural activity was observed during mismatch after CT training, but not after NT training. No increase to playback halt is observed in either condition, in line with the recordings from Attinger et al. shown for comparison in 4B. The model also reproduced recovery of mismatch responses observed by Attinger et al: Subsequently training the model originally trained in the NT paradigm by exposing it to the CT conditions installed an increase in neural activity under the mismatch condition 4C and 4D). One source of difference between the simulations and experimental results is the simple nature of the neuron model.

**Figure 4.**
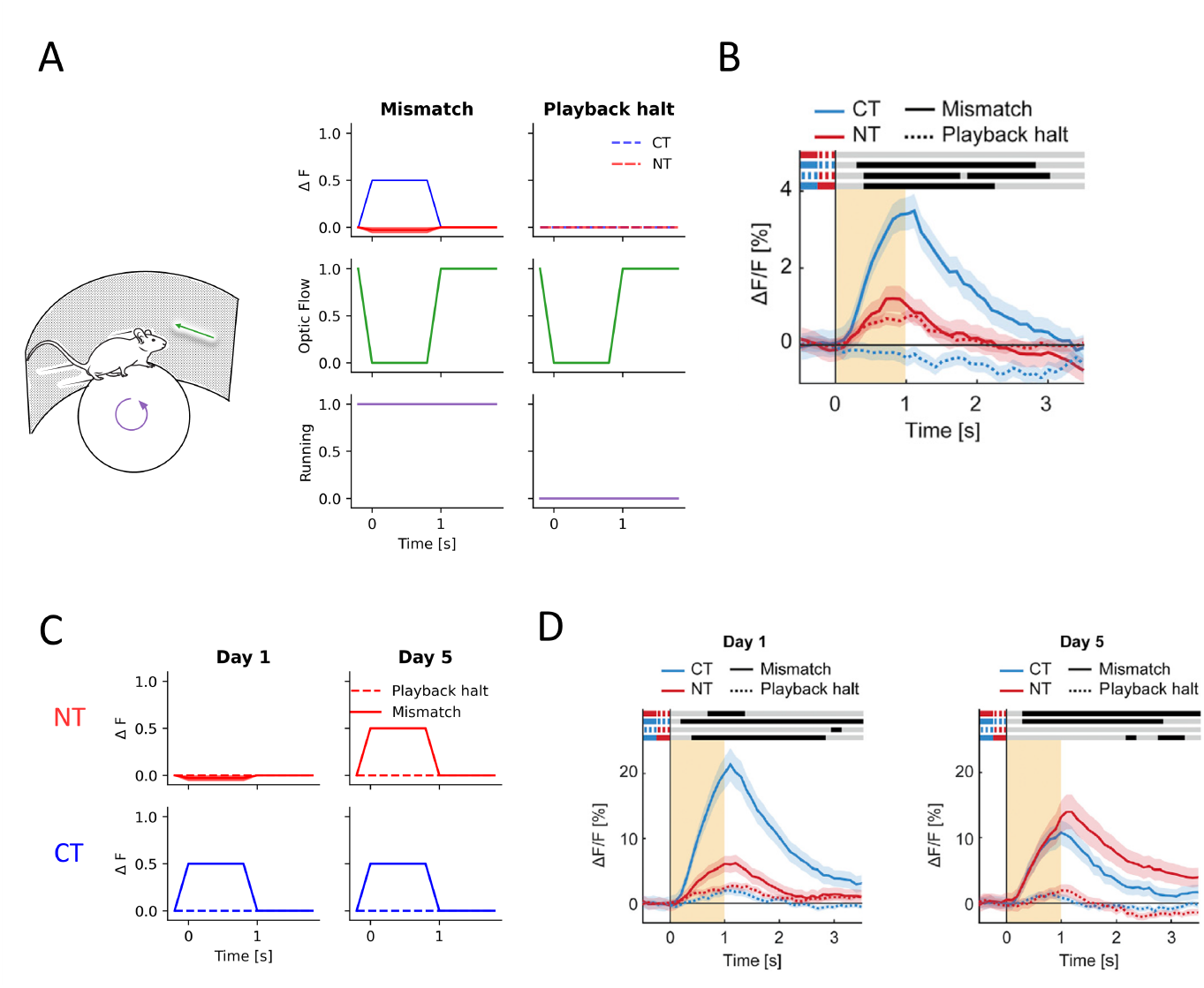
The proposed microcircuit reproduces experimentally observed mismatch responses. (A) Left: VR setup with optic flow stimuli (green) independently controllable of motor state (purple). Right: Response of model V1 error neurons in the model to mismatch and playback halt conditions defined by sudden halt of optic flow from *t* = 0*s* to *t* = 1*s* while running (mismatch) and standing still (playback halt). After experiencing coupled training (CT), i.e. contingent sensorimotor experience, a mismatch response was observed. This was not the case after non-coupled training (NT) with visual feedback uncorrelated to locomotion. Error bands (partially invisible) indicate standard deviation across five random seeds influencing the sampling of the training sequences. (B) Calcium imaging responses to the two conditions in mice V1. Delta F/F (in somatic Ca^2+^) on the vertical axis is a reflection of neural firing rate. The orange area indicates duration of visual halt, shading indicates SEM. (C) Responses of model error neurons after training in CT and NT conditions (left, “day 1”) and after the model has been subsequently trained in the coupled paradigm (right, “day 5”). Note the appearance of the mismatch response in the red continuous curve. (D) Observed neural responses in mice initially trained in the CT and NT conditions, before and after subsequent coupled training. Panels B and D were reproduced from Attinger et al., 2017, published in Cell, Copyright Elsevier.

Since no receptor dynamics are modeled and the experimental data is a low-pass filtered Ca^2+^-response, the rise of activity is faster in the model. Without an explicit model of somatic Ca^2+^, the activity decay is instantaneous in the model, whereas the recordings in Figure 4B, D show a slow signal decay characteristic for calcium imaging.

### 3.2 The higher-level visual area reduces prediction errors

After positive evaluation of the microcircuit, we tested its capacity to reduce prediction errors when integrated into a hierarchical predictive coding model of visual processing (see section 2.2). In the pretraining phase illustrated in the left panel of Fig. 5A, we restricted predictions about optic flow to top-down feedback from the motor area to V1-EN. As shown in Fig. 5B, coupled training minimized prediction errors in this phase much more efficiently than non-coupled training. Here, we used the Animal dataset described in section 2.2.2. In the non-coupled paradigm, mean squared error (MSE) as measured by input to error neurons amounted to 3.19±0.00 (a.u.) after this phase, compared to 0.74±0.00 in the coupled condition. This correlates well with the experimentally observed necessity of coupled training to truthfully detect sensorimotor mismatch (cf. Fig. 4B). While in the Animal dataset used here the lateral movement of the background image created homogeneous optic flow patterns, we also tested learning of multiple non-homogeneous flow patterns. As shown in Suppl. Fig. 11, the parallel microcircuits from motor neurons to V1-EN (Suppl. Fig. 10) were able to accurately learn the corresponding forward models.

**Figure 5.**
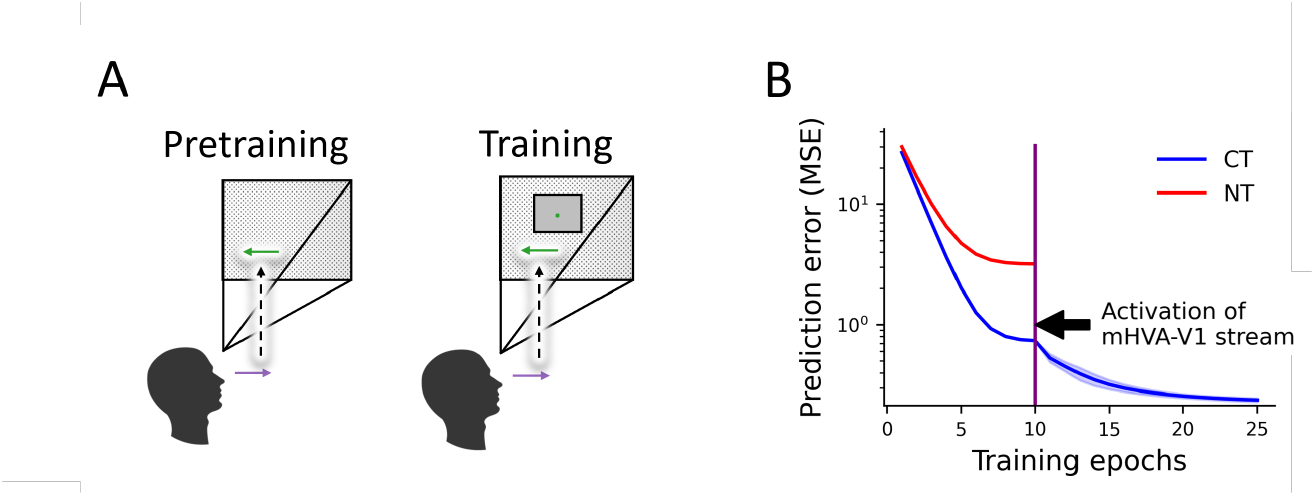
Learning to predict external and internal causes of optic flow. (A) Pretraining setup: the model is trained to predict the consequences of its own movements in an empty environment. Training setup: objects are introduced, that cast a static image onto the retina due to the observer moving along them, while the background patterns move across the retina. As in previous figures, the colored vectors refer to absolute/world-centric movement (purple) and pattern movement relative to the observer’s retina (green). (B) Pretraining (epoch 1-10) reduces prediction errors most effectively if conducted in the coupled paradigm (CT) as opposed to the non-coupled paradigm (NT) both introduced in Fig. 2B and in the main text. Activation and training of the second visual stream from mHVA to V1 after epoch ten further reduces prediction error. Error bands (too small to be visible in epoch 1-10) indicate one standard deviation computed across four runs with randomly initialized weights.

After training the motor-to-visual stream, we then tested in the training phase whether addition of the second visual stream (mHVA to V1) provided a reduction in prediction errors quantified by activity in V1 prediction error neurons. To do so, the visual-to-visual stream of the model was activated and the weights of the pretrained motor-to-visual stream were frozen, putatively corresponding to a halt of critical period plasticity. In Fig. 5B, this moment is indicated by the vertical line. Training then proceeded until convergence of the error signal which took approximately 15 epochs (each epoch consisting of a single iteration through the ten sequences of the dataset). This further reduced the MSE from 0.74 *±* 0.00 to 0.24 *±* 0.01. Qualitatively, the effect of feedback from area mHVA to V1-EN can be understood by the following scenario: With the motor-to-visual stream trained on background-only inputs (i.e. the pretraining paradigm described in 2.2.2), and before activation of the second visual stream, an external object moving independently from the background elicited an increase in nPE activity (Fig. 6A, second column). After activating and training the mHVA-V1 stream, predictions from model area mHVA successfully minimized prediction errors (Fig. 6A, third column), i.e. optic flow including the moving animal was better predicted. This can be explained by representation of the object-generated optic flow patterns in model area mHVA that is then fed back as top-down prediction to V1 where the full bottom-up optic flow is represented, cancelling the object-generated prediction errors there (cf. Eq 8). While separate training of both streams proved to be the most effective strategy, we demonstrated that in principle also both streams can be trained simultaneously (Supplementary Material 7.4).

**Figure 6.**
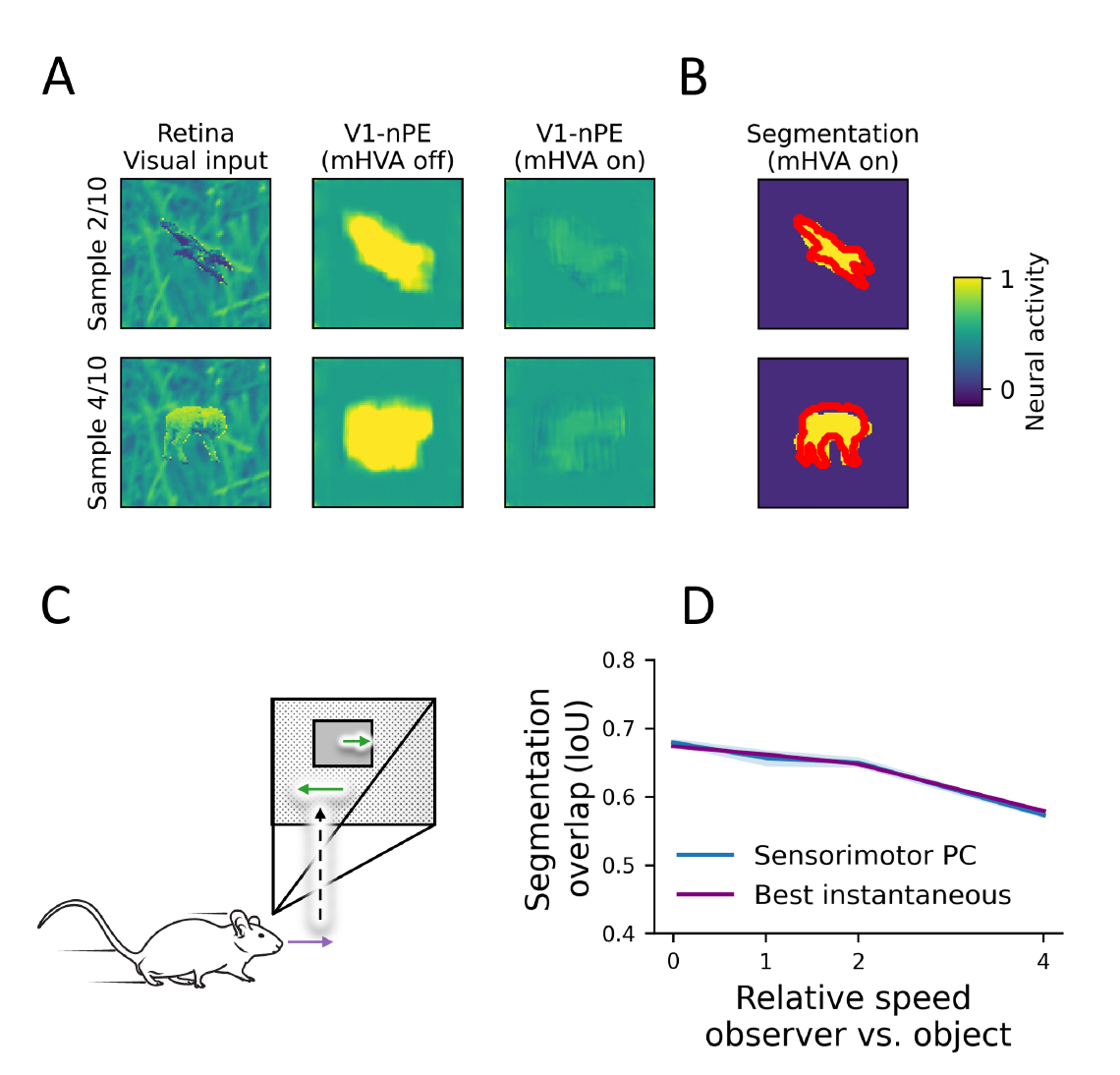
Segmentation of external causes through visual feedback. (A) Same setting as in Figure 5. First column: Retinal inputs showing an eagle and a lamb against a similarly colored background. Second column: A model with the motor-to-visual stream trained in the coupled paradigm correctly minimizes prediction errors in the background, but the external object moving slower than expected elicits activity in nPEs. pPEs are not shown for simplicity but show the inverse pattern. Third column: after activating and training feedback from mHVA to V1, prediction errors are further minimized. (B) Outlining the fully trained model’s guess about the spatial extent of the external object, the segmentation mask obtained from top-down signals from model area mHVA to V1 is shown in yellow over the ground truth outlined in red. (C) Non-zero relative speed of the object relative to the observer. In contrast to the stable perception paradigm, the object image moves across the retina (upper green arrow). (D) Segmentation performance under varying relative speeds between object and observer during observer movement. Prior to evaluation, the two-stream model was trained in the paradigm of C on the Animal dataset. The error bands indicate one standard deviation, calculated across four randomly initialized runs. The baseline in purple is the optimum at the time step of evaluation (see section 2.3).

### 3.3 Visual predictions segment external causes

If, as hypothesized above, model area mHVA indeed represents external causes of optic flow that cannot be explained by self-motion, it should be possible to use its activity to segment moving objects from the background. We obtained segmentation masks from the model as described in section 2.3. The model accurately captured the outline of external objects as shown in Fig. 6B.

We evaluated segmentation performance of the model on objects moving relative to both the observer and background. To do so, the Animal dataset was modified: In addition to background motion correlating with the movement state of the agent, the external object now moved at an independent speed in the opposite direction (Fig. 6C). We found that the model was capable of shifting the segmentation mask across retinal space as indicated by the high IoU scores (introduced in section 2.3) in Fig. 6D. Here we selected appropriate timepoints for evaluating readout accuracy to ensure that the object was still in the model’s field of view: For speeds [0, 1, 2, 4] these were [9, 9, 6, 4] frames into the sequence. Across relative movement speeds, model performance was on par with the baseline derived by optimal segmentation of the instantaneous optic flow (described in more detail in section 2.3), and both mildly decreased as expected with faster movement speed.

### 3.4 Higher visual areas encode object identity

For the visual system to guide behavior, it does not suffice to know where an externally moving object is located (shown by the correct placement of the segmentation masks); also its identity needs to be inferred. Evaluating the information content of model area mHVA in this context also allows us to better understand its computational role.

To read out representational content from the model, we used the modified FashionMNIST dataset described in section 2.2.2. First, both generative streams were trained according to the procedure described in section 3.2. As before, the binary locomotion state was coupled to movement of the background pattern across the retina, while the object’s image remained stable in the center. In this phase, use of 100 stimuli (10 per class) proved sufficient. Then linear decoding was used to estimate readout accuracy of object class as described in section 2.4. The resulting classification accuracy of 81.6% on unseen (test) images in area mHVA and 82.3% in mHVA2 shows class-specificity (Tab. 1) and decent generalization to unseen data. Succeeding rejection of the null hypothesis (‘all populations have the same readout accuracy’) using a Welch’s ANOVA, we performed a Games-Howell post-hoc test to determine the significance of the pairwise differences between the populations (for a detailed description of the statistical methods see section 7.7).

**Table 1:** Model area mHVA allows readout of object class. Percentage of correctly classified stimuli evaluated on the train/test split of the ten classes of the modified FashionMNIST dataset as described in the main text. Rows denote the cell population: mHVA, mHVA2, V1-Direction Selective (DS) cells, full retinal input as a baseline, and chance level. The standard deviation is calculated across four randomly initialized runs.

| Area | Acc. train | Acc. test |
| --- | --- | --- |
| mHVA | $94.0 \pm 0.2$ | $81.6 \pm 0.7$ |
| mHVA2 | $93.3 \pm 0.7$ | $82.3 \pm 0.5$ |
| V1-DS | $88.8 \pm 0.3$ | $79.6 \pm 0.4$ |
| Full vis. | $92.7 \pm 0.2$ | $76.8 \pm 0.1$ |
| Chance | 10.0 | 10.0 |

Showing an increase in decodability compared to lower areas, readout accuracy on test data was slightly but significantly higher in area mHVA (81.6%) than in V1-DS cells (79.6%) with *p <* 0.03, and also significantly higher than the baseline obtained from directly reading out the raw retinal activity patterns (76.8%, *p <* 2.5e-3). The difference between areas mHVA2 and mHVA was not significant. Lastly, in all areas, the drop between readout accuracy on training and test data suggested overfitting of the readout classifier. We conclude that learning optic flow patterns drives learning of behaviorally useful and generalized representations and emphasize that more class-specific representations are likely to be acquired via additional learning mechanisms in the ventral stream. Purely reconstructing activity patterns in previous areas did in our case not improve representation readout, suggesting the need for additional mechanisms such as explicitly enforcing sparsity Dora et al., 2021 or an external feedback (reward or supervision). Due to the invariance of optic-flow to changes in luminance, the approach employed here can be expected to be more robust to changes in lighting condition than approaches that rely on texture, i.e. patterns in static luminosity.

### 3.5 Unexpected events elicit elevated neuronal responses

To further examine the general conjecture that model area mHVA codes for optic flow caused by externally moving objects, we recorded the neural dynamics of the network in two conditions. In a shape-from-motion paradigm illustrated in Fig. 7A, the external object from the Animal dataset began to move in front of the background, followed by movement of the observer so that the object image appeared statically on the retina. In the model trained in the coupled-training paradigm, this movement onset elicited an activity increase in mHVA neurons with receptive fields on the object, but not for those with receptive fields on the background (Fig. 7C, bottom row). In V1-PE neurons with receptive fields on the object, but not outside, a strong transient response was observed, reminiscent of retinotopically organized error coding Zmarz and Keller, 2016. Due to the suppressive effect of area mHVA on V1-PEs (see 5), the rapid decay in activity was most likely caused by top-down feedback. We note that the “stable perception” paradigm in which the observer maintains the object image statically on the retina was chosen for consistency with the previously described simulations and was not necessary to elicit the described neural responses (object movement sufficed).

**Figure 7.**
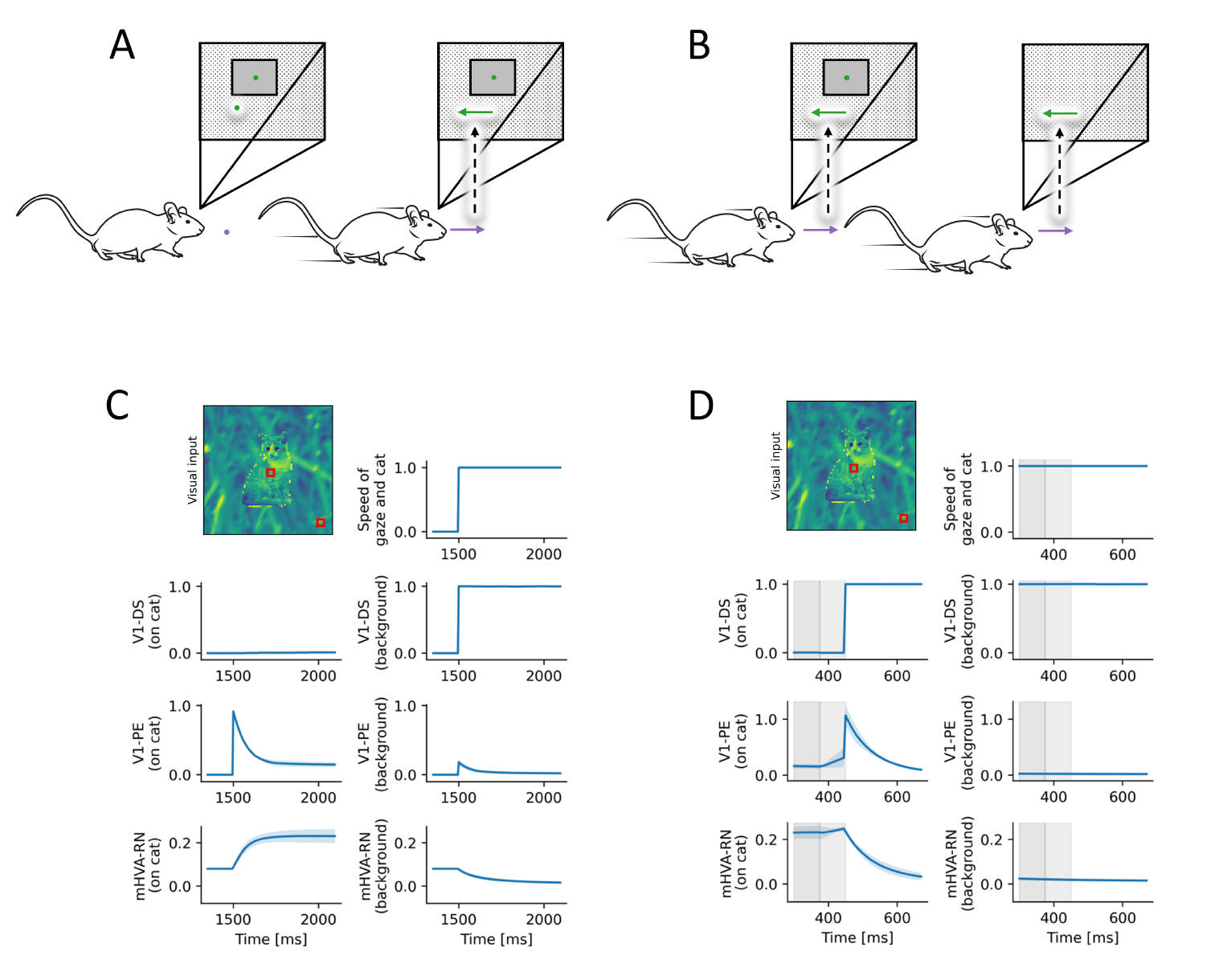
Neuronal dynamics during movement onset and object removal. (A) Illustration of movement onset. Green arrows indicate movement of patterns across the retina, the purple arrow movement of the observer. Left: the starting point is static, symbolized by points in green (relative to the retina) and purple (world-centric). Right: both the object and the observer begin to move, so that the object appears static on the retina. (B) Object removal: after the observer moves synchronously with it (left), the object suddenly disappears (right). (C) Neuronal activity during movement onset as illustrated in A. Responses were recorded across two sites indicated by red squares. The light blue band (invisible) indicates one standard deviation calculated across four randomly initialized runs. (D) Neuronal activity during the object removal outlined in B. At the end of the area shaded in grey, the cat image was suddenly removed, where the lightly shaded area indicates a transition time of one frame during which optic flow estimation is disturbed until a reliable estimate could again be made. Representations in area mHVA take a number of time steps to decay (bottom row). This difference between prediction and presence results in a ‘spike’ of error signals in V1 (third row).

To confirm that the response in area mHVA was truly linked to object presence, we recorded the model’s response to unexpected object disappearance (Fig. 7B). The network with two streams trained (CT) on the Animal dataset was first moving to maintain an image from the Animal dataset statically on the retina. After a fixed amount of time steps, the image was suddenly removed (illustrated in Fig. 7B). Following object removal, a non-instantaneous decrease of mHVA activity was observed for neurons with receptive fields on the object. Furthermore, we again observed a spike of activity in V1-PE neurons with receptive fields on the object (Fig. 7D), fitting well with the response increase observed in Zmarz and Keller, 2016.

In this setting, responses can thus be explained by model area mHVA representing the distortions in optic flow caused by the external object. Of the higher visual areas in mice, area PM has been suggested to code for external objects in a similar fashion Andermann et al., 2011. Sudden removal then creates a divergence between the still present prediction of optic flow-distortion by a moving object and the sensed optic flow that is now completely explained by egomotion. Consequently, mHVA reduces its activity, reflecting the novel absence of external optic-flow distortions.

## 4 Discussion

### 4.1 Summary of results

We developed a scalable microcircuit with an analytically tractable learning rule to learn the visual consequences of egomotion (Fig. 2). The microcircuit replicated sensorimotor mismatch signals observed in V1 of mice under various conditions and in an experience-dependent manner (Fig. 4). Subsequently, we integrated the microcircuit into a model designed for inferring the causes of optic flow (Fig. 3). To our knowledge, this joint generative model is the first to functionally incorporate motor-to-sensory microcircuits into a biological model of visual inference, conforming to hierarchical predictive coding. This extended model successfully learned to predict self-generated optic flow patterns across the visual field (Fig. 5B), and was capable of learning multiple non-homogeneous optic flow patterns (Supplementary Fig. 11). The benefit of the added visual-to-visual stream was firstly demonstrated by its capacity to reduce prediction errors in V1 more effectively than a pure motor-to-visual model when external objects were introduced (Fig. 5A). We then showed that this efficiency in error-minimization was due to top-down signalling of a segmentation mask by model area mHVA, highlighting the external object. Lastly, we confirmed that area mHVA not only learned the 2D shape of the object, but that population activity of mHVA neurons also contains enough information for accurate readout of object class.

### 4.2 The neural circuitry of predictive coding

While the predictive coding framework offers a compelling account of inference and learning in perception, its empirical predictions are still under scrutiny (Green et al., 2023; Leinweber et al., 2017; Pennartz et al., 2019; Walsh et al., 2020). With regard to sensorimotor mismatch, the question is to what extent neuronal mismatch signals can be explained purely by locomotion-induced gain as argued by Muzzu and Saleem, 2021, 2023. A recent case in favor of true generative motor-to-visual feedback was made by Vasilevskaya et al., 2023. Together with the alignment with experimental results, use of a biologically plausible learning rule and its strong computational benefits (object representation learning without external supervision, optic flow-based segmentation), the predictive coding account for sensorimotor mismatch appears compelling.

Another point of debate is the necessity of dedicated error neurons. Although recent evidence suggests prediction error-coding by SST interneurons in the posterior parietal cortex of mice (Green et al., 2023), dedicated error neurons are not required per se for prediction-based inference and learning. Indeed, Mikulasch et al., 2023 developed a model of hierarchical (unimodal) predictive coding with error computation in basal dendrites (see also Urbanczik and Senn, 2014). Due to the algorithmic similarity of how bottom-up sensory inputs are compared to top-down predictions in their model compared to ours (since both are derived from Rao and Ballard, 1999), many aspects proposed here, such as the learning rule, functional modularization and its mapping to brain areas are translatable to an implementation with dendritic error computation.

It should be mentioned that sensorimotor mismatch is not a purely cortical phenomenon and has also been observed in the cerebellum (Hull, 2020, see also Lisberger, 1988). There, however, mismatch responses are mostly linked to learning of sensorimotor transformations and motor control (Albus, 1971; Hull, 2020; Ito, 1970; Lisberger, 1988; Marr, 1969; Stone & Lisberger, 1986) without object segmentation, whereas the circuits modeled here underlie a perceptual function: the parsing of visual inputs in a dynamic world. It thus appears that the brain developed the same principle of comparing the predicted outcome of movement to the observed state at least twice for different purposes.

### 4.3 Translation to primate visual perception

Predictive processing is often postulated as a general organization principle across species (Friston, 2005). How then does the model translate to primate perception? Rodents and primates share patterns of cortical organization that have been hypothesized to serve predictive processing (Bastos et al., 2012; Brooks & Cullen, 2019). Recordings of motor-induced activity changes in visual areas (Gutteling et al., 2015; Haarmeier et al., 1997; Lubinus et al., 2022; Paradiso et al., 2019) support the existence of a motor-to-visual stream as in the model, underlining the subjective experience of motor-to-visual coupling reported in Pelah and Barlow, 1996. Nevertheless, the location of sensorimotor mismatch computation may well lay in extrastriate areas (Brooks & Cullen, 2019; Parker et al., 2020), as primate V1 appears to be less affected by movements than in mice (Talluri et al., 2023).

Regarding visual-to-visual feedback, model area mHVA maps well onto the primate middle temporal cortex (MT), based on the shared functional properties of direction-selectivity (differently weighted inputs from pPE and nPE; (Albright, 1984; Maunsell & Van Essen, 1983b)), large receptive fields, strong feedback connections to early visual areas (Clavagnier et al., 2004), and its functional role in perceiving structure from motion (Andersen et al., 1996; Born & Bradley, 2005; Buračas & Albright, 1997; Duncan et al., 2000; Grunewald et al., 2002; Handa et al., 2008)). Computationally, this mapping aligns well with the interpretation of Born and Bradley, 2005 who concludes from a review of passive recording studies that “[a]ll together the evidence rather strongly suggests that MT neurons are critically involved in segmenting an image into separately moving parts”. Following our model architecture, we go a step further by postulating that this is achieved through neurons tuned to world-centric as opposed to retinocentric movement. Such *real motion cells* were indeed found in MT (Erickson & Thier, 1991), further supported by Pitzalis et al., 2020 and Sulpizio et al. (manuscript in preparation). While the flow-parsing theory of Warren and Rushton, 2009 explains such real motion coding in MT with a subtraction operation *within* MT (bottom-up received optic flow minus self-generated optic flow), our model allocates only the outcome to MT and the subtraction operation to error neurons in lower visual areas. Here, our model provides a novel perspective on the large number of feedback connections from MT to lower visual areas (Maunsell & Van Essen, 1983a) as part of a larger generative model employed in the interpretation of sensed optic flow.

### 4.4 Related work

Based on psychophysical evidence of object movement perception during selfmotion, Royden and Holloway, 2014 constructed a model to identify the outlines of moving objects. This was achieved through comparison of bottom-up optic flow patterns to a memory bank of optic flow templates. In contrast to our model, however, no neural mechanisms for the learning of the templates, nor for the wiring of the comparison operation were implemented.

Within the framework of predictive coding, most models typically operate purely in the visual domain, whether static (Rao & Ballard, 1999), or dynamic (Brucklacher et al., 2023; Lotter et al., 2020). Extensions of predictive coding that take into account the multimodality of sensory processing in the brain were proposed by Keller and Mrsic-Flogel, 2018; Pennartz, 2015, 2022 in theory and implemented e.g. by Pearson et al., 2021. Another type of multimodal generative model is learning of motor-to-visual forward models based on corollary discharges, wherein motor commands are taken to constitute a non-sensory modality. On a neural level, such models have been proposed by Hertäg and Sprekeler, 2020; Mikulasch et al., 2022, in contrast to our model without the ability to account for externally-caused optic flow. Interestingly, Hertäg and Sprekeler, 2020 demonstrated that signed prediction error responses emerge under visual feedback contingent with motor state in a biologically detailed model, and investigate the role of the involved interneurons. The microcircuit developed in section 2.1 can thus be seen as a useful abstraction over the interneuron-level circuits developed by Hertäg et al., lending it more easily to functionally powerful computations and the abovementioned integration with a visual-to-visual stream. In contrast to Mikulasch et al., 2022 and Hertäg and Sprekeler, 2020, the used datasets go beyond one-dimensional visual inputs and the model is capable of solving visual tasks such as figure-ground segmentation and, to a limited degree, classification. To our knowledge, the model put forward here is the first to integrate motor-to-sensory forward models and sensory-sensory predictive coding within a functional model of generative visual perception.

### 4.5 Limitations in performance

Segmentation performance (Fig. 6B and D) is naturally dependent on the accuracy of the underlying optic flow-extraction through V1 direction-selective cells. Here, accuracy can be assumed to correlate positively with the resolution of the input signal. Based on the relatively low resolution used due to limitations in computational resources (40 x 40, 80 x 80 pixels per frame), we expect significant improvements when using input sequences of higher resolution (and thus wider networks). Another factor that would significantly improve segmentation is the use of depth information through motion parallax or - especially in primates - stereovision. An interesting extension of the current model would thus be a joint generative model of optic flow and depth for 3D segmentation and recognition from binocular inputs. Depth is also an informative cue for segmentation of partially overlapping objects moving at the same speed - whereas the current model depends on distinct speeds. Lastly, it should be stressed that self-generated optic flow is by no means irrelevant (Gibson, 1950), but provides an important reafference signal to inform motor execution. Interestingly, as discussed in section 4.2, such closed-loop control also relies on sensorimotor prediction errors (Albus, 1971).

### 4.6 Conclusion

In summary, this study presents a computational model that seamlessly integrates motor-to-sensory microcircuits into a hierarchical predictive coding framework for visual perception. Notably, the model’s use of generative feedback for external object segmentation provides a novel angle to the ongoing discourse regarding the functional importance of top-down connections to early sensory areas in neural networks.

## 5 Acknowledgments

We thank Sander Bohte, Valentina Sulpizio and Thede Witschel for helpful discussions.

This project has received funding from the European Union’s Horizon 2020 Framework Programme for Research and Innovation under the Specific Grant Agreement No. 945539 (Human Brain Project SGA3 to G.P., F.M. and C.P.). This project has received funding from the European Research Council under the Grant Agreement No. 820213 (ThinkAhead), the PNRR MUR projects PE0000013-FAIR, and IR0000011–EBRAINS-Italy to G.P.

A research stay connected to this project was funded for M.B. through the NENS Exchange Grant program of the Federation of European Neuroscience Societies (FENS).

## 6 Data availability

The Python code to reproduce the results of this paper can be found at: https://github.com/matthias-brucklacher/LearningMotorFeedback

## 7 Supplementary material

### 7.1 Loss function and convergence of learning rule

To derive a principled loss function corresponding to the learning rule of Eq. 4, we first consider *w*_−_ and then use the symmetry of the circuit to generalize to *w*_+_. Plugging the firing rate of the nPE neuron from Eq. 1 into the weight update yields

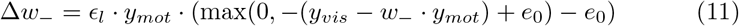

Here we assume the model to be moving (*g* = 1), otherwise no learning would take place as can be seen from Eq. 4. In the coupled training paradigm, it holds true that

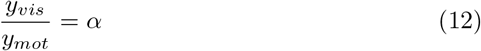

with *α* ∈ ℝ. The resulting weight update is then the rectified linear function shown in blue in Fig. 8 with zero value derived from Eq. 11 at

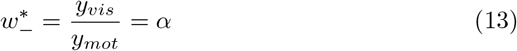

**Figure 8.**
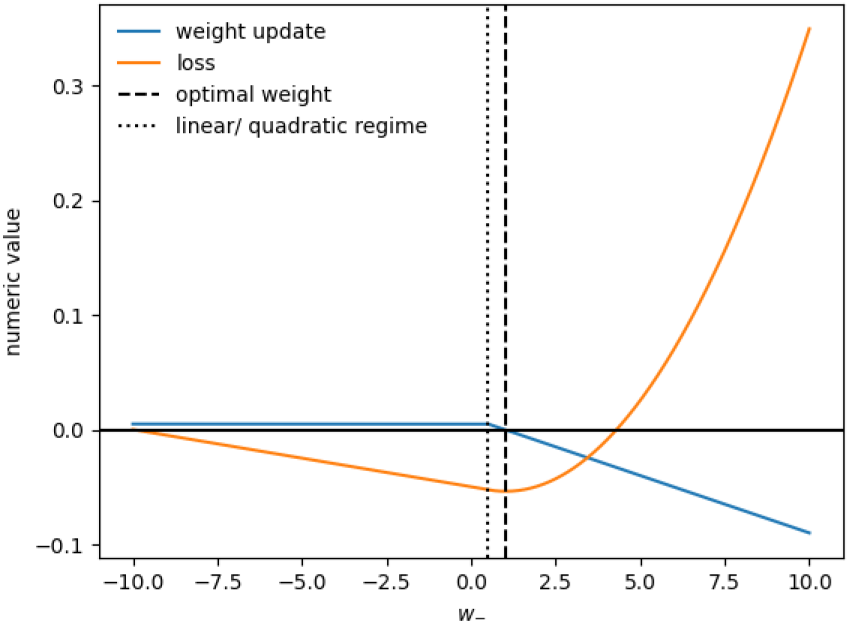
Loss function underlying the learning rule. As can be seen from the global minimum of the loss function (Eq. 15) depicted in orange, the weight updates (blue) are guaranteed to drive the weight towards the optimum value 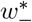 (dashed vertical line).

At this value, the net input to the error neuron is zero. Integrating weight updates yields the (negative) loss function on which learning performs gradient descent, shown in orange in Fig. 8. Based on the constant and negatively sloped linear parts of its negative derivative shown in blue, it can be seen that its global minimum lies at the optimal weight 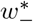 (denoted by the dashed line in Fig. 8) with strictly monotonic increase in both directions. This guarantees convergence of the learning rule to 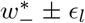. The transition from a linear to a quadratic regime (dotted line in Fig. 8) occurs at the firing threshold of the error neuron,where the net input to the neuron becomes zero

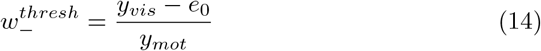

Integration of the weight update in each regime and division by the learning rate then yields the loss

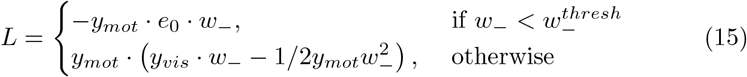

with integration constants appropriately chosen (integration from *w*_−_ = 0). Analogous derivation for the pPE neurons yields the optimal weight

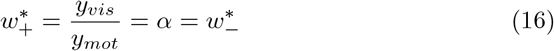

### 7.2 Construction of the Animal dataset

To construct the Animal dataset, ten images of animals where obtained from pixhttps://pixabay.com under the Pixabay content license allowing free use and modification of the images for commercial and non-commercial use https://pixabay.com/service/terms/. As an illustration, one image can be found under https://pixabay.com/de/photos/pferd-grau-konik-weide-l%C3%A4uft-2388274/, and links to all images are listed in data/original/image credits.txt in the GitHub repository). The animal images were cropped out of their original background, grayscaled, and resized so that the longer edge was 50 pixels long. Then, the images were pasted in front of the grass background texture (80 x 80 pixels) obtained from https://www.patternpictures.com/full-frame-green-grass-texture/ under the Pattern Picture license allowing unrestricted use (https://www.patternpictures.com/license/) allowing lateral shifting according to movement of the observer and the object (as described in section 2.2.2).

### 7.3 Hyperparameters of the two-stream model

The used hyperparameters are listed in Tab. 2.

**Table 2:** Hyperparameters of the two-stream model.

| Symbol | Parameter | Value |
| --- | --- | --- |
| $c_{thresh}$ | Segmentation threshold | 0.71 |
| $e_0$ | Baseline activity V1-EN | 0.5 a.u. |
| $\epsilon_{inf}$ | Inference rate mHVA & mHVA2 | 0.05 |
| $\epsilon_l$ | Learning rate pretraining | 200 |
| $\epsilon_l$ | Learning rate training | 0.02 |
| - | mHVA feature maps | 4 |
| - | mHVA2 feature maps | 4 |
| kernel_size | Kernel size mHVA-V1 | [4, 4] |
| - | Kernel size mHVA2-mHVA | [4, 4] |
| - | Convolutional stride mHVA-V1 | 1 |
| - | Convolutional stride mHVA2-mHVA | 1 |

**Table 3:** Full report on the outcome of the Welch’s ANOVA for the readout of object class. ddof1: degrees of freedom (numerator), ddof2: degrees of freedom (denominator), p-unc: uncorrected p-values, np2: Partial eta-square effect sizes. This figure supports main text section 3.4.

|  | Source | ddof1 | ddof2 | F | p-unc | np2 |
| --- | --- | --- | --- | --- | --- | --- |
| 0 | Population | 3 | 5.698583 | 142.910062 | 0.000009 | 0.953495 |

**Table 4:** Full report of the post-hoc pairwise test (Games-Howell). Columns A and B refer to the neuronal populations tested in a pairwise manner. Columns: mean: average readout accuracy on the test data across the randomly initialized runs, diff: difference between mean values, se: standard error, df: adjusted degrees of freedom, pval: Games-Howell corrected p-values, hedges: Hedges effect size.

| Games-Howell post-hoc |  |  |  |  |  |  |  |  |  |  |
| --- | --- | --- | --- | --- | --- | --- | --- | --- | --- | --- |
|  | A | B | mean(A) | mean(B) | diff | se | T | df | pval | hedges |
| 0 | Full vis. | MST | 0.76775 | 0.82275 | -0.05500 | 0.003075 | -17.883614 | 3.497725 | 0.000481 | -10.996195 |
| 1 | Full vis. | MT | 0.76775 | 0.81550 | -0.04775 | 0.003955 | -12.071843 | 3.292599 | 0.002467 | -7.422680 |
| 2 | Full vis. | V1-DS | 0.76775 | 0.79625 | -0.02850 | 0.002407 | -11.842487 | 3.846641 | 0.001247 | -7.281654 |
| 3 | MST | MT | 0.82275 | 0.81550 | 0.00725 | 0.004863 | 1.490940 | 5.615486 | 0.499418 | 0.916743 |
| 4 | MST | V1-DS | 0.82275 | 0.79625 | 0.02650 | 0.003714 | 7.135718 | 5.603904 | 0.002063 | 4.387578 |
| 5 | MT | V1-DS | 0.81550 | 0.79625 | 0.01925 | 0.004470 | 4.306674 | 4.825987 | 0.029702 | 2.648068 |

### 7.4 Simultaneous learning of visual and visuomotor stream

In the main results, motor-to-visual and visual-to-visual streams were trained independently. Upon activation and training of the mHVA-V1 stream, the connections in the motor-to-visual stream were frozen. In the development of cortico-cortical brain connectivity (Price et al., 2006), not much is known about the developmental order of feedback connections from higher visual areas to V1 relative to connections from motor areas to V1. In macaques, hints come from the work of Baldwin et al., 2012 that suggests the presence of feedback connections from area MT to V1 about two weeks after birth. To investigate sensitivity of the model to separate training phases, we studied simultaneous training of all plastic connections. To investigate this, we combined both pretraining (background only) and training (Animals) data into an interleaved dataset. Here, one in ten sequences contained an animal, the other nine were empty. Inference was conducted as described in section 2.2 and plasticity was enabled in both streams. In general we found this training paradigm to be quite unstable compared to the separate training phases. To achieve segmentation performance, several modifications proved helpful:

1. A high learning rate in the motor-to-visual stream compared to the visual-to-visual stream. This enforced learning of a correct motor-to-visual stream on a fast timescale and thus partially decoupled the two learning problems.
2. A large ratio (*>*10:1) of empty sequences over sequences with objects in the training data to ground the motor-to-visual stream.
3. Adapting the segmentation threshold. We found predictions of the visual-to-visual stream to be about 30% weaker in this joint training paradigm compared to the separate training paradigm. Lowering the threshold (of summed up predictions) for assigning a point in space to an external object prevented the segmented areas from becoming too small.

With these tweaks, the jointly trained model reached an IoU of 0.56 compared to 0.68 of the separately trained model. The worse performance illustrated in Fig. 9 was likely due to noisier predictions resulting from interactions between the two streams via V1 error neurons.

**Figure 9.**
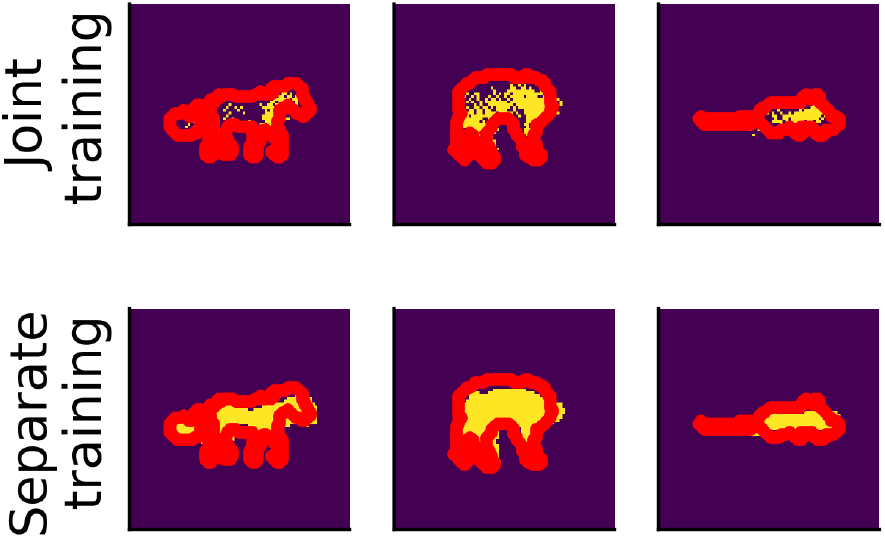
Joint training leads to noisier segmentation performance compared to separate training of both streams. Segmentation mask from mHVA→ V1-feedback (yellow) over the ground truth of the object outline (red) on the Animal dataset (cf. Fig. 6). Top row: model trained with plasticity enabled in both streams, bottom row: separate pretraining and training phases as described in section 2.2.2

**Figure 10.**
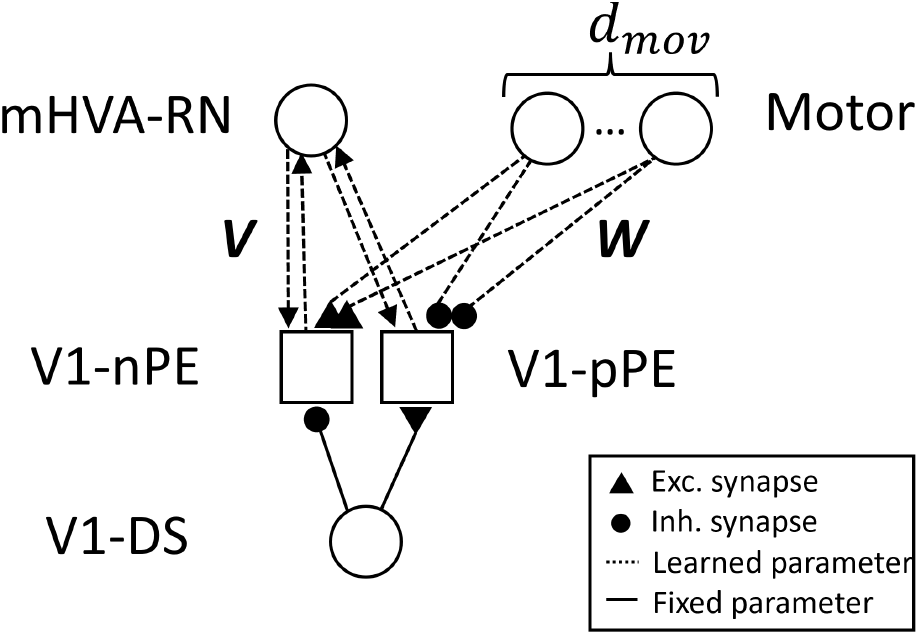
Forward models from multidimensional action states. In an extension of the model shown in Figure 3, multiple motor neurons encode a *d*_*mov*_-dimensional action space. For abbreviations see Fig. 3.

**Figure 11.**
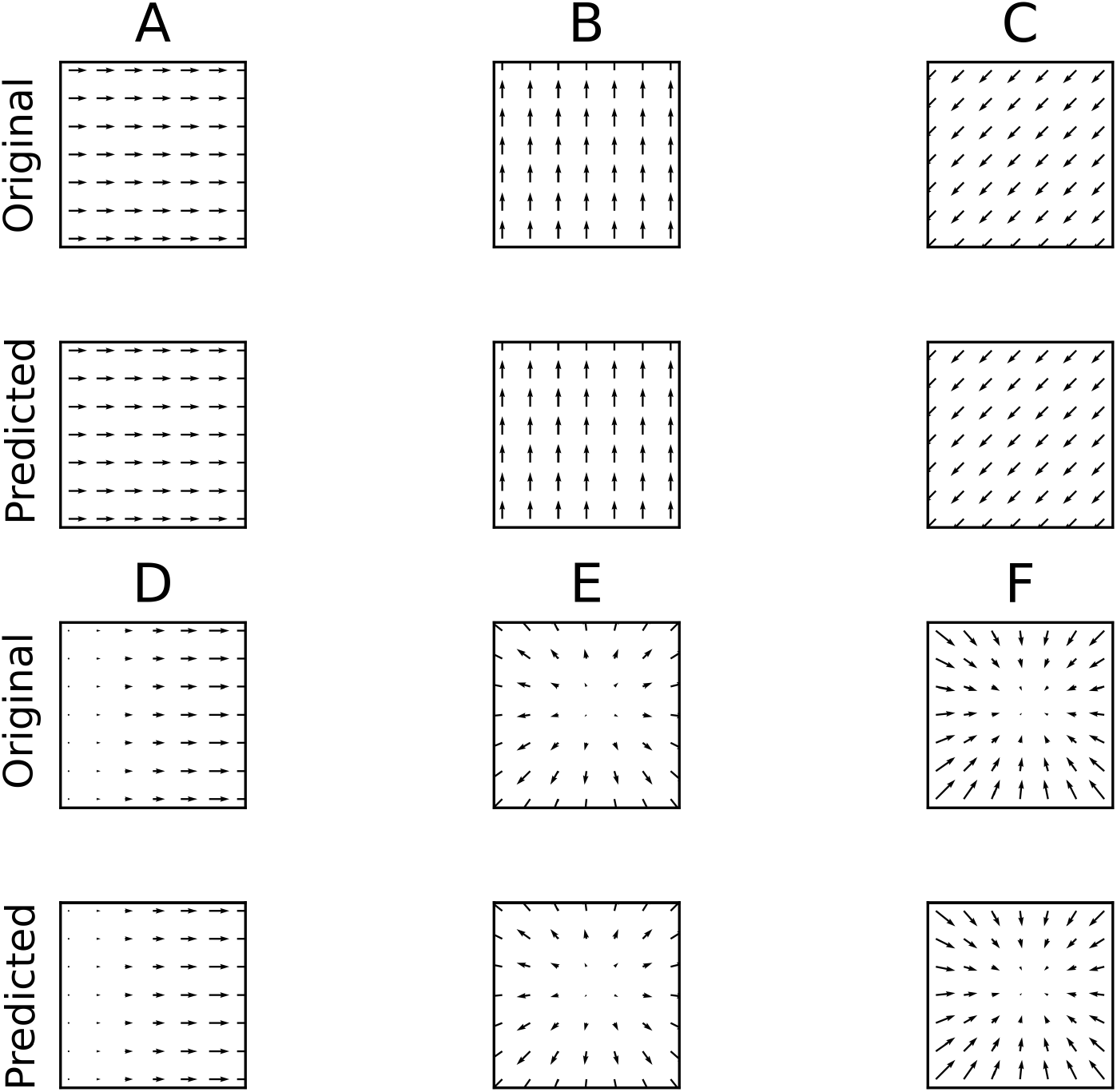
Learning outcomes of multiple movements in one model. Optic flow input patterns of the six movement dimensions are shown above predictions from motor stream to V1 after training. “Original” optic flow is obtained from V1-DS neurons and predicted optic flow from mtr-V1 signals (averaged across V1-nPE and V1-pPE at each point in retinotopic space). The vectors (plotted at every fourth pixel column/row) are computed by subtracting opposing signal components: the *x*-component of the vector is given by rightward minus leftward (predicted) optic flow, the *y*-component by upwards minus downwards (predicted) optic flow. The actions corresponding to the chosen optic flow patterns are (A) turning leftwards, (B) looking downwards, (C) looking up and to the right, (D) following a leftward curve while walking, (E) moving forward and (D) moving backwards.

### 7.5 Learning distinct forward models from multiple actions

Distinct movements create differing optic flow patterns, such as when walking forward as opposed to turning around. The model’s capacity to learn such mappings was demonstrated by training it on the multidimensional dataset of artificially generated optic flows described in section 2.2.2. Figure 10 illustrates the wiring of the model. After training the motor-to-visual stream for five epochs, the model’s prediction from the motor stream closely matched the original optic flow patterns, as shown in Fig. 11.

### 7.6 Addition of a second higher visual area

Vision is generally understood as a hierarchical cortical process (Lee & Mumford, 2003)). Also most neural networks models of generative perception are strictly hierarchically organized (Dayan et al., 1995; Dora et al., 2021; Rao & Ballard, 1999), although a strict hierarchy is debated (St-Yves et al., 2023; Suzuki et al., 2023) as implemented e.g. in (Salvatori et al., 2022). In rodents, the hierarchy of motion-sensitive areas is not mapped out well yet, but in general processing appears to be hierarchically organized as well (Siegle et al., 2021). Inspired by this evidence for hierarchical processing, we extended our model in depth by adding an area on top of mHVA (Fig. 12). In primates, where the hierarchy is better understood, this newly introduced model area roughly maps onto the medial superior temporal cortex (MST) because it is further removed from thalamic sensory input in the visual hierarchy than MT (roughly corresponding to mHVA in the model) (Born & Bradley, 2005; Felleman & Van Essen, 1991). Interconnected with mHVA neurons via linear error neurons (mHVA-EN, for simplicity not split into positive and negative counterparts), area mHVA2 learns a generative model of activity patterns in area mHVA. Connections are convolutional with stride 1, kernel size 4 and mapping to four feature maps in area MST. With the input resolution of the modified FashionMNIST dataset described in section 2.2.2, the number of neurons in mHVA2 is thus 34 x 34 x 4 = 4,624.

**Figure 12.**
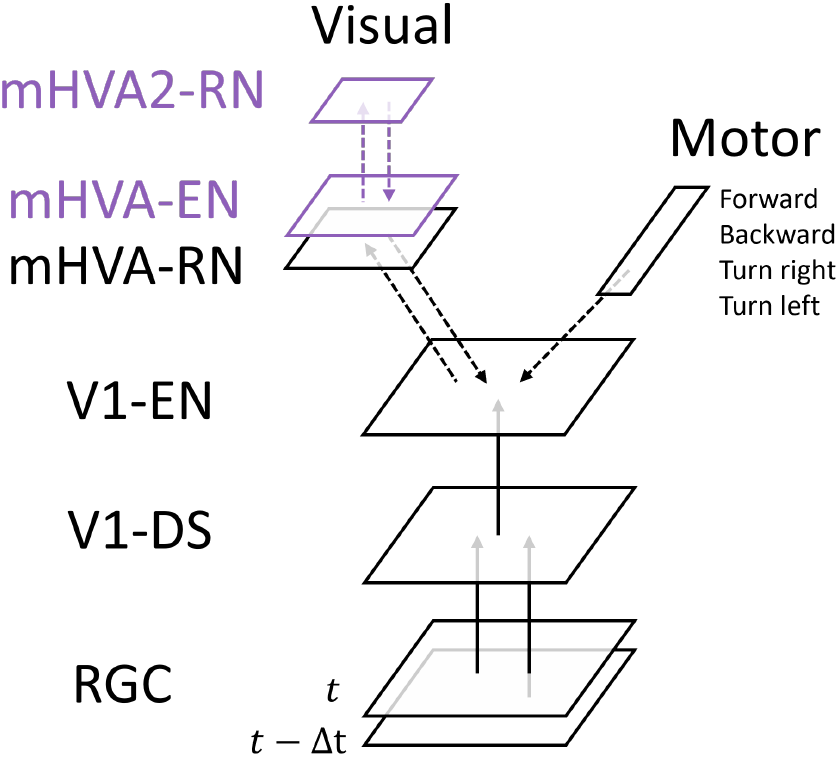
Extension of the visual-to-visual stream. Shown in purple is the added model area mHVA2 and its connection to area mHVA via an added population of error neurons (mHVA-EN). Compare to Fig. 3.

As in area mHVA (see section 2.2), inference and learning in area mHVA2 are driven by error signals from the area below. Since we were interested in the representations formed in mHVA2, we did not implement an influence of mHVA-EN on area mHVA-RN as common in other predictive coding implementations and Eq. 6 remains unchanged. Thus, the remaining network function is not affected by the addition of area mHVA2. Training and readout evaluation then progressed as described in sections 2.2.2 (training) and 3.4.

### 7.7 Statistical methods

To analyze whether significant differences between readout accuracy (on the test dataset) across network areas were present, we first conducted a Welch ANOVA using the pingouin package in Python (https://pingouinstats.org/build/html/index.html). The null hypothesis (i.e., no significant difference in readout accuracy across representation neurons in the three network areas) was rejected with *p <* 1e-5 (Tab. 7.7).

Subsequently, multiple pairwise comparisons were conducted using the *pingouin* implementation of the Games-Howell post-hoc test, which led to the corrected *p*-values reported in section 3.4 of the main text. All comparisons are shown in Tab. 7.7.

